# Theta Oscillations Organize Spiking Activity in Higher-Order Visual Thalamus during Sustained Attention

**DOI:** 10.1101/219626

**Authors:** Chunxiu Yu, Iain M. Stitt, Yuhui Li, Zhe Charles Zhou, Kristin K. Sellers, Flavio Frohlich

## Abstract

Higher-order visual thalamus plays a fundamental but poorly understood role in attention-demanding tasks. To investigate how neuronal dynamics in higher-order visual thalamus are modulated by sustained attention, we performed multichannel electrophysiological recordings in the lateral posterior-pulvinar complex (LP/pulvinar) in the ferret (*Mustela putorius furo*). We recorded single unit activity and local field potential during the performance of the 5-choice serial reaction time task (5-CSRTT) that is used in both humans and animals as an assay of sustained attention. We found that half of the units exhibited an increasing firing rate during the delay period before stimulus onset (attention-modulated units). In contrast, the non-attention-modulated units responded to the stimulus, but not during the delay period. Spike-field coherence of only the attention-modulated neurons significantly increased from the start of the delay period until screen touch, predominantly in the theta frequency band. In addition, theta power and theta-gamma phase-amplitude coupling were elevated throughout the delay period. Our findings suggest that the theta oscillation plays a central role in orchestrating thalamic signaling during sustained attention.

**Significance:** Impaired sustained attention can be deadly, as illustrated by the number of motor vehicle accidents that are caused by drivers not reacting quickly enough to unexpected events on the road. Understanding how electrical signaling in higher-order visual nuclei, such as the LP/pulvinar, is modulated during tasks that require sustained attention is an important step in achieving a mechanistic understanding of sustained attention, which will eventually lead to new strategies to prevent and treat impairment in sustained attention.

## Introduction

Sustained attention is defined as the allocation of processing resources to rare but important events during prolonged periods of time [1]. Sustained attention is a key element of attention models [2] and integrates multiple behavioral processes including executive control for controlling competing impulses and executing planned actions to appropriate stimuli [3].

In humans, sustained attention is typically measured with the continuous performance task (CPT), which requires the participant to respond to an infrequent stimulus over prolonged time. Neuroimaging studies support an active role of the fronto-parietal attention network in sustained attention [4] but subcortical structures have also been proposed to be part of the network substrate of sustained attention [5]. Sustained attention is not only impaired in attention deficit hyperactivity disorder (ADHD) [6-8] but also depression [9], bipolar disorder [10], and schizophrenia [11]. Interestingly, impairment of sustained attention persists in patients with bipolar disorder [12] and major depressive disorder [13] even after achieving remission.

In animal model species, one task that has been extensively used to delineate the substrate of sustained attention is the five-choice serial reaction time task (5-CSRTT) that probes spatial sustained attention [14]. In the 5-CSRTT, animals respond to a visual stimulus at one of five stimulus locations after a delay period. The 5-CSRTT is a widely used task that has provided fundamental insights into the cellular and molecular mechanisms of sustained attention, and is used for the evaluation of candidate compounds for the treatment of ADHD [15].

Despite the extensive study of sustained attention at the behavioral level in both humans and animal models, surprisingly little is known about the circuit dynamics of sustained attention. Recently, we reported that the 5-CSRTT engaged synchrony of oscillatory activity in the frontoparietal network of the ferret [16, 17]. Phase synchronization both at the microscale of neuronal action potentials and the mesoscale of the population synaptic activity reflected in the local field potential was elevated in the theta-frequency range.

In contrast, the role of subcortical structures in oscillatory network interactions during sustained attention has remained mostly unknown. The thalamus plays a major role in regulating the thalamo-cortical network dynamics that reflect vigilance levels [18, 19]. Since vigilance requires sustained attention, we asked if higher-order thalamus is engaged in the 5-CSRTT. In particular, we investigated how spiking and oscillatory network activity is modulated during the delay period during which the animals pay attention to the potential stimulus locations.

## Materials and Methods

Three spayed female ferrets (*Mustela putorius furo*, 4 months of age at the beginning of the experiments) were trained to reach satisfactory task performance and subsequently implanted with electrode arrays in the LP/Pulvinar complex for combining neurophysiological measurements with behavioral data collection. All procedures were approved by the Institutional Animal Care and Use Committee of the University of North Carolina at Chapel Hill and followed National Institutes of Health guidelines for the care and use of laboratory animals.

### Behavioral apparatus

Ferrets were trained in a touch-screen based version of the 5-Choice Serial Reaction Time Task (5-CSRTT) [20]. Training took place in an enclosed and sound-attenuated custom-built operant chamber (51 x 61 x 61 cm^3^). The chamber consisted of a touch screen monitor (Acer T232HL 23-inch touch screen LCD display) covered with a black Plexiglass mask. The mask exhibited five equally sized square cut-outs (windows, 7 x 7 cm), in one of which the stimulus was presented in each trial. Below the openings, there was a shelf on which the animals could comfortably rest their front legs when nose-poking the touchscreen. The opposite wall was equipped with a lick spout centrally positioned 6 cm above the floor for water delivery combined with an infrared (IR)-based proximity detector and an LED light. The behavioral chamber also included a “house light” mounted on the ceiling of the chamber opposite the monitor to provide feedback for correct and incorrect trials and a speaker (HP Compact 2.0 speaker) to deliver auditory tones. Infrared videography was performed during all sessions (Microsoft LifeCam Cinema 720p HD Webcam with filter that blocks infrared light removed). The entire behavioral setup was controlled by a data acquisition device (USB 6212, National Instruments, Austin, TX) and custom-written Matlab scripts (Mathworks, Natick, MA) that used functionality from the Psychophysics toolbox [21] for precise temporal control of stimulus presentation.

### 5-choice serial reaction time task (5-CSRTT)

In the 5-CSRTT [20], the animals initiated a trial by approaching the lick-spout at the rear end of the behavior chamber. Upon initiation, a 5 sec delay period started during which no stimuli were presented and the ferrets had to prepare and sustain attention to the five windows (cut-outs in the mask on the touchscreen). After the delay ended, a white solid square (stimulus) was randomly presented for 2 sec in one of five windows. For correct trials, touching the stimulus window during the 2 sec of stimulus presentation or in the first 2 sec after the stimulus was turned off (hold period, HP) triggered a 0.5 sec tone, illumination of the lick spout light, a water reward release at the lick spout (Fig.1A, left). For incorrect trials, touching a window before the stimulus onset (PreTouch) or touching an adjacent incorrect window after 5 sec delay (MissTouch), or failing to respond to the stimulus at all (NoTouch, omission) caused a 1 sec white noise auditory stimulus, illumination of the house light, and a subsequent 6 sec time-out period, in which no water was delivered (Fig. 1A, right). Following the 8 sec inter-trial interval after correct responses or 6 sec time-out period after incorrect responses, the lick-spout light turned on to indicate the availability of the next trial. The session was terminated after 60 trials or 40 min, whichever came first. Criteria of >80% accurate stimulus detection and <20% omission for at least five consecutive sessions were used during the training.

**Figure 1:**
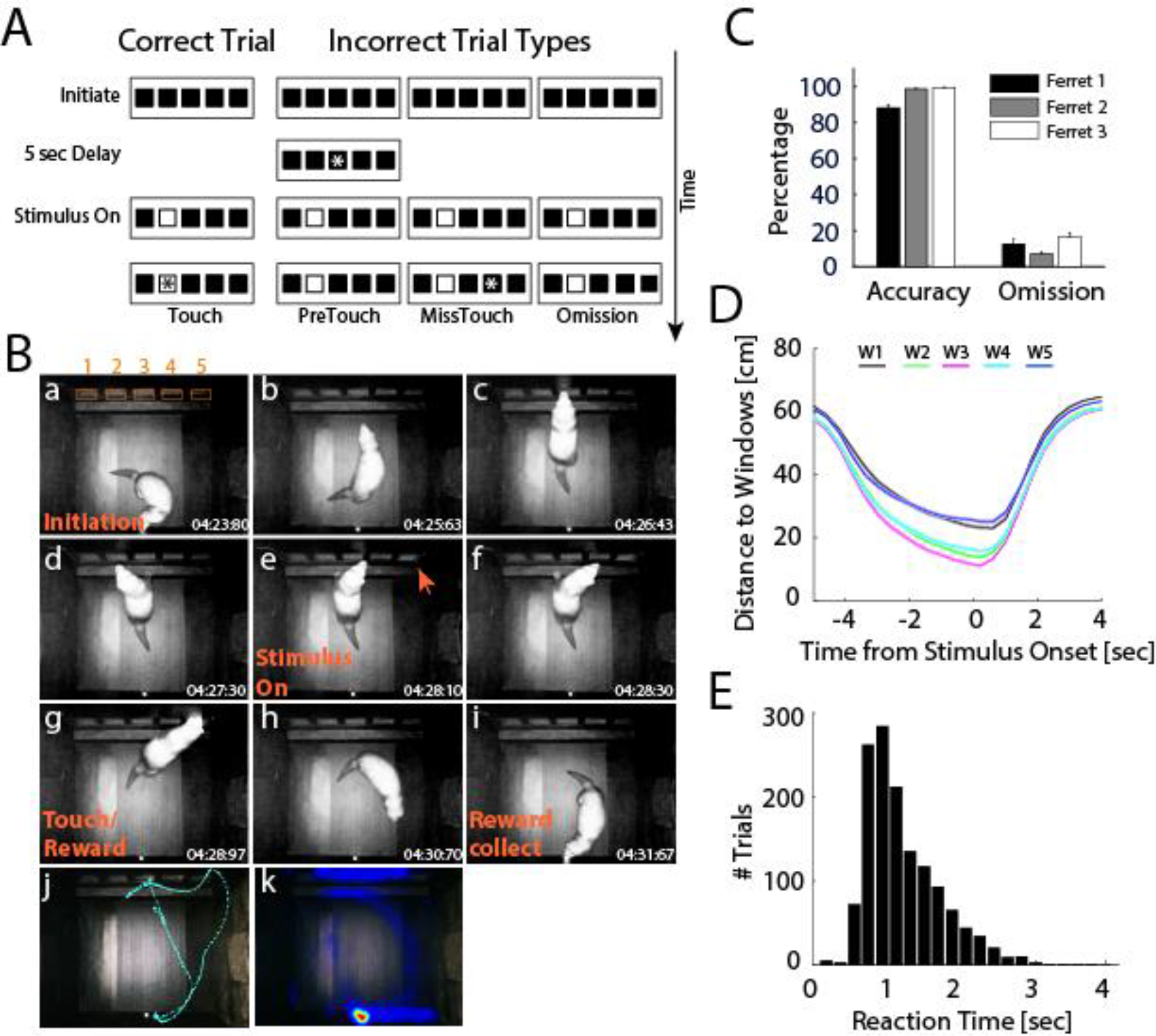
Sustained Attention Task in Freely-Moving Ferrets. **A** Illustration of trial sequences of the 5-CSRTT. Each trial begins with illumination of the water spout, which is centrally placed on the back wall of the chamber. The ferret initiates the trial by approaching the water spout, which is equipped with an infrared proximity sensor. Then, the spout light is extinguished and the 5 sec delay period starts during which the animal is required to sustain attention to the five windows on the front wall of the chamber. A white solid square (stimulus) will randomly present in one of five windows after the delay ends. Nose-poke to the stimulus window during stimulus presentation (2 sec) or in the first 2 sec after stimulus offset (hold period, HP) triggers a tone and delivery of water reward at the spout (correct trial, left). Touch by the ferret is indicated with an asterisk. If the ferret responds before the stimulus (PreTouch), or touches an incorrect window (MissTouch), or fails to respond before the end of the HP (Omission), a 6 sec time-out (TO) period is introduced where the house light is on and no water is delivered (right). After collecting the reward (8 sec time window) or at the end of the TO period, a new trial can be started. **B** Representative photographs of one animal during a single trial: a, initiation; b, turn to the stimulus windows (numbered 1 through 5); c-d, paws on the platform and checking for occurrence of stimulus; e, stimulus on; f, find the stimulus; g, nose poke the stimulus window; h, turn back to collect the reward; I, collect the reward; j, complete trajectory for a single trial obtained from video tracking. K, heatmap of animal locations during the session. Time stamps are shown in the corner of each frame. **C** Behavioral performance. Mean accuracy and omission rates across sessions. Error bars: Standard error of the mean (SEM) across sessions. **D** Mean distance between animal location and stimulus location as a function of time for correct trials. The shorter distances to middle windows (W2 – W4) indicate that animals were centered relative to the stimulus windows before stimulus onset. **E** Distribution of touch reaction times for correct trials. In most trials, the correct window was touched during stimulus presentation (RT < 2 sec).

### Behavioral training

Ferrets were water deprived and received free access to water (60ml) each day after training and testing. Body weight was monitored daily and never dropped below 85% which would have triggered discontinuation of water deprivation by protocol. The ferrets underwent a multistep training plan that included five subsequent training levels: 1, arena habituation 2, touch-reward association 3, single stimulus-reward association 4, task initiation 5, 5-CSRTT. Following this training schedule, the ferrets were successively advanced to the final task, in which the ferrets were required to initiate the trial by nose-poking the water spout. Ferrets were trained and tested daily on a 5 days on / 2 days off schedule.

### Surgery

After successful training, the ferrets were surgically implanted with a chronic multichannel electrode array (2x8 tungsten electrode array, 250μm spacing, low-impedance reference electrode with the same length on the same array, MicroProbes Inc, Gaithersburg, MD) in LP/Pulvinar. Surgical procedure was similar to those previously described [16, 22]. Briefly, ferrets were first injected with ketamine/xylazine (30 mg/kg of ketamine, 1-2 mg/kg of xylazine, IM) for anesthesia induction, and then were intubated and anesthetized with inhaled isoflurane (1.5-2%) in 100% medical oxygen (mechanically ventilated, 10-11 cc volume, 50 bpm). Body temperature was maintained between 38-39°C with a water heating blanket. Electrocardiogram, end-tidal CO2 and partial oxygen saturation were monitored throughout surgery. The skull was exposed and a craniotomy was performed over the target area (centered at 13 mm anterior to lambda, 3.6 mm lateral from midline) [23]. A small slit was made into the dura before insertion of electrode array (7.4 mm from cortical surface). Electrode arrays were secured in place with dental cement and several non-penetrating skull screws. A separate ground wire was implanted in cortex of the same hemisphere. The wound margins were sutured together and anesthesia was reversed. Ferrets received standard postoperative care with 3 days of meloxicam for pain relief (0.2 mg/kg, IM) and 7 days of clavamox to prevent infection (12.5-13mg/kg, oral).

### Electrophysiological recording

The animals were given two weeks of recovery time before they were retrained on the final task to again reach stable performance (typically a few sessions). At this point, wireless electrophysiological recordings were performed with a MCS 2100 system (Multichannel Systems, Reutlingen, Germany). Broadband data (1 Hz to 5 kHz) were collected (sampling rate: 20 kHz) and digitally stored for offline analysis. Simultaneously with acquisition of the neuronal data, time-locked acquisition of relevant behavioral events (trial initiation, screen touch, reward retrieval) was performed through digital input channels on the wireless recording system. Behavioral data were stored separately by a custom-written Matlab script. In addition, high-resolution infrared videography (30 frames/s) was performed that was synchronized to the neuronal data acquisition.

### Histology

To verify electrode locations in the LP/pulvinar complex, electrolytic lesions were induced at the completion of recording by passing current through the four electrodes (two middle and two corner) of the microelectrode array (5 μA, 10s, unipolar). The damage visible in the sections stem from this lesioning protocol. Animals were then humanely euthanized with an overdose of sodium pentobarbital (IV injection) and perfused with 0.1M PBS and 4% paraformaldehyde solution in 0.1M PBS. The brains were post-fixed and cut in 60 μm coronal sections using a cryostat (CM3050S, Leica Microsystems), and then stained for cytochrome oxidase [24]. Imaging was acquired using a widefield microscope (Nikon Eclipse 80i, Nikon Instruments, Melville, NY). Electrodes outside of LP/pulvinar were excluded from analysis.

### Data analysis

Behavioral data were analyzed with custom-written Matlab scripts. The main measurements of interest were: (1) identity of the window touched, (2) reaction time after visual stimulus onset, and (3) animal location relative to windows on touchscreen. Equally, MissTouch, NoTouch and PreTouch trials were also tracked. Performance accuracy in detecting the stimulus was defined as percentage of the number of correct responses divided by the total number of correct and incorrect (MissTouch) responses. The video recordings were analyzed offline with Ethovision XT (Noldus, Leesburg, VA) and the three-point animal-tracking module to detect the animal location (sample rate 30 frames/s). Heatmaps were generated for demonstration of animal trajectories. We also manually reviewed video recordings and coded the orientation of the animal immediately prior to the stimulus onset. Only correct trials with the animal orienting to the screen at the time of stimulus onset were included in further analysis.

Electrophysiology data were analyzed by a combination of custom-written Matlab scripts. Trials exhibiting simultaneous large amplitude deflections across channels (artifacts in electrophysiological recordings) were excluded. Single units were extract with the Plexon Offline Sorter (Plexon Inc, Dallas, TX) spike sorting software. Briefly, multiunit activity was detected by high-pass filtering the raw continuous data with a Butterworth 2nd order filter with 300Hz cut-off. Action potentials were detected by threshold crossing (-4 times the standard deviation of the high-pass filtered signal). Spikes were identified, collected and sorted using the T-distribution expectation maximization algorithm [25] and manual inspection with the Plexon Offline Sorter. Spikes with shorter than 1 msec refractory period were removed. Neural activity within a window of -7 to 5 sec aligned on stimulus onset was averaged in 200-msec bins and averaged across trials to construct the peri-stimulus time histogram (PSTH). The PSTHs were then z-scored by subtracting the mean baseline firing rate (measured from a baseline window from -2 to 0 second before trial initiation) and dividing by the standard deviation. To classify “attention-modulated” vs. “non-attention-modulated” units, neural activity within 2 second before the stimulus onset was compared to a baseline window from -2 to 0 second before the trial initiation (two-tailed t test, p<0.05). The 2 sec window was chosen based on visual inspection of the behavioral video. The ferrets usually began facing the windows and remained in an oriented posture around 2 second before stimulus onset.

To investigate the rhythmic structure of LP/pulvinar activity, local field potentials (LFP) were computed by low-pass filtering the raw continuous data with a Butterworth 2nd order filter with 300Hz cut-off. To study the synchrony between LP/Pulv spikes and the LFP at different epochs during the task, we computed the spike-field coherence of simultaneously recorded spike trains and LFP using multi-taper analysis provided by the Chronux data analysis toolbox [26]. The spike-field coherence was calculated in 2 sec windows using the *‘coherencycpt’* function for each epoch (before initiation, before stimulus onset, before screen touch and after screen touch). We corrected coherence values to remove the effects of unequal number of trials in each session [26, 27]. The corrected spike-field coherence was computed using the formula of T (f) = atanh(C (f))-1/ (df-2), where C(f) is the raw coherence value; df is the degrees of freedom; df= 2*K*N, where K is the number of tapers and N is the number of trials. The corrected coherence estimates were pooled across the population. Two-way ANOVA was used to compare the modulation of the coherence at different epochs during the task.

Cross-frequency phase-amplitude coupling (PAC) was computed to assess the degree with which high frequency oscillations are temporally organized by the phase of low frequency oscillations in the LP/Pulvinar. PAC was defined as the phase locking value (PLV) between low frequency signals and amplitude fluctuations of high frequency signals occurring at the lower carrier frequency. First, an LFP signal *x*(*t*) was convolved with a low frequency Morlet wavelet with carrier frequency *f*_1_

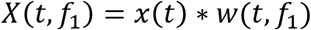

where * denotes the convolution operation. Then the analytic amplitude of the same signal *x*(*t*) at a higher carrier frequency *f*_2_ was convolved with a wavelet with carrier frequency *f*_1_

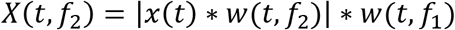

The instantaneous phase of each time series was then computed by taking the argument of the real and imaginary components of the time frequency estimates. Finally, PAC is defined as the PLV between low and high frequency signal components

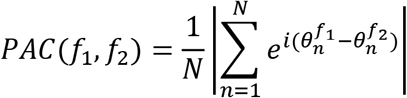

PAC values were bound between 0 and 1, with 0 indicating no relationship, and 1 perfect alignment of high frequency amplitude modulations with low frequency phase. To examine PAC extensively in frequency-frequency parameter space, we computed PAC between low frequencies (*f*_1_) ranging from 1 to 10 Hz in 0.25 Hz steps, and high frequencies (*f*_2_) from 10 to 80 Hz in 1 Hz steps. This analysis revealed prominent PAC between theta and gamma frequencies in LP/Pulvinar. After identifying this frequency-frequency band of interest, we repeated analysis using a phase preserving 4^th^ order Butterworth filter and Hilbert transform instead of Morlet wavelet convolution to maximally capture time frequency fluctuations in the theta/gamma band of interest (theta 3-6 Hz, gamma 30-60 Hz). Finally, to investigate how theta/gamma PAC is modulated by the 5-CSRTT, we computed PAC in sliding windows of 200 msec (step size 100 msec) aligned to stimulus onset.

## Results

### Behavioral Performance

To probe the role of LP/Pulvinar neuronal dynamics in sustained attention, we trained 3 ferrets to perform a five choice serial reaction time task (5-CSRTT, Fig. 1A-B). The 5-CSRTT was designed as a task in which freely moving animals make a choice in each trial to get a reward. Trials were self-initiated at a lick spout in the back of the behavioral apparatus and a touch-screen with five stimulus locations was used to display the targets and capture the behavioral responses. All ferrets reached criterion performance across recording sessions with high accuracy (Fig. 1C, Mean ± SEM, Ferret 1: 88.1% ± 1.85%, Ferret 2: 98.7% ± 0.38%, Ferret 3: 99.4% ± 0.29%) and low omission rates (Ferret 1: 12.6 ± 2.77%, Ferret 2: 7.10% ± 1.40%, Ferret 3: 16.7% ± 2.12%). Video tracking data confirmed that animals moved towards the screen during the delay period (Fig. 1D), with animals reaching their final position close to the screen approximately 2 seconds before stimulus onset. This implies that ferrets allocated attention to the stimulus windows in anticipation of the visual stimulus during this time period. In addition, the reaction time distribution for correct trials indicated that animals responded quickly to stimuli, with most responses within 2 seconds of stimulus onset (Fig. 1E). We then asked if neuronal firing in the LP/Pulvinar was modulated during the delay period.

### Single Unit Activity in LP/Pulvinar

We implanted microelectrode arrays into LP-Pulvinar and confirmed implantation locations through histological reconstruction of the recording sites (Fig. 2A). We found that about half of the LP/Pulvinar single neurons (n = 130/259 units) displayed a progressively increasing firing rate during the delay period of the task (Fig. 2B-D). Since no stimuli were present in any of the windows during the delay period, such elevated firing rates suggest that the excitability of this subpopulation of LP/Pulvinar neurons is modulated by behavioral or mental engagement of the animal with the possible locations of the future stimulus (referred to as attention-modulated). The remaining 50% of neurons (n = 129/259) did not exhibit significant changes in firing rate during the delay period. However, both attention-modulated and non-attention-modulated LP/Pulvinar neurons displayed increases in firing rate in response to the visual stimulus. This increase was significantly stronger for attention-modulated neurons (Unpaired t test, p <0.001). Thus the delay period of the 5-CSRTT led to a ramping up of spiking activity in a subpopulation of LP/Pulvinar neurons.

**Figure 2:**
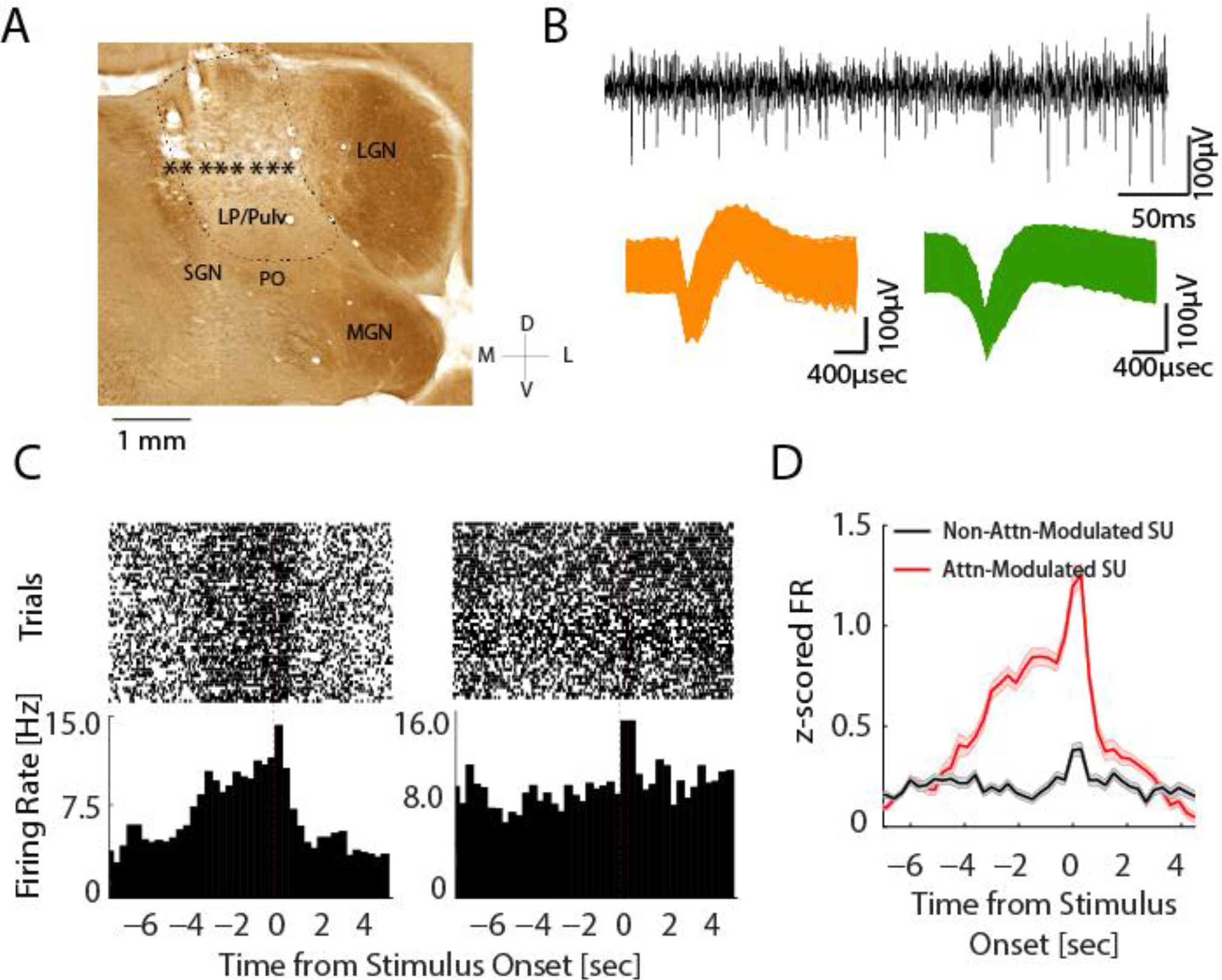
Single-Unit Responses during Task Performance. **A** Representative coronal section; stars indicate the estimated locations of the electrode tips. Electrodes outside of LP/pulvinar (LP/Pulv) were excluded. LGN, lateral geniculate nucleus; MGN, medial geniculate nucleus; PO, nucleus of the posterior group; SGN, suprageniculate nucleus. **B** Top: Example of high-pass filtered raw trace from representative recording session. Bottom: Action potentials of two representative single units. **C** Peri-event raster plots (top) and peri-event histograms of the corresponding firing rates (bottom) of two representative neurons. The unit on the left exhibited an increasing firing during the delay period, whereas the unit on the right did not show such modulation but rather increased its firing rate after stimulus onset. **D** Z-score normalized population firing rate of attention-modulated (red, n=130) and non-attention-modulated units (black, n=129). Shaded areas indicate SEM. Attention-modulated units gradually increased their firing rate during the delay period. Both neuron types displayed a transient increase in firing rate after stimulus onset, with stronger responses in attention-modulated neurons. Shaded areas indicate standard error of the mean (SEM).

### Modulation of Spike-Field Coherence in the Theta Band

We next asked if the task-related activity of the neurons in LP/Pulvinar exhibited a mesoscale organization that would be reflected in the LFP. To answer this question, we computed spike-field coherence (SFC) of both attention-modulated and non-modulated neuron subpopulations during different epochs of trials. We found SFC in the theta band (∽5 Hz) throughout task performance (Fig. 3). Strikingly, SFC in the theta band differed for attention-modulated and non-modulated neurons. A two-way ANOVA with neuronal response type (“attention-modulated” or “non-modulated”) and task segments (before initiation, before stimulus onset, before touch and after touch) as factors revealed a significant main effect of response type (F_1, 1035_ = 55.8, p < 0.001), main effect of timing (F 3,1035 = 4.04, p = 0.007), and significant interaction between these factors (F_3_, 1035 = 4.07, p = 0.007). Post-hoc comparison showed that SFC in the theta frequency band was significantly higher for attention-modulated neurons than for non-modulated neurons during all trial epochs (p < 0.05), with the exception of the period after the screen-touch (p > 0.05). While no changes in theta SFC was observed across task epochs for non-modulated neurons, theta SFC was significantly higher before touch than before trial initiation for attention-modulated neurons (p < 0.05). These results suggest that neurons whose activity fluctuated with attention selectively synchronized to theta rhythms based on behavioral demands throughout the task.

**Figure 3:**
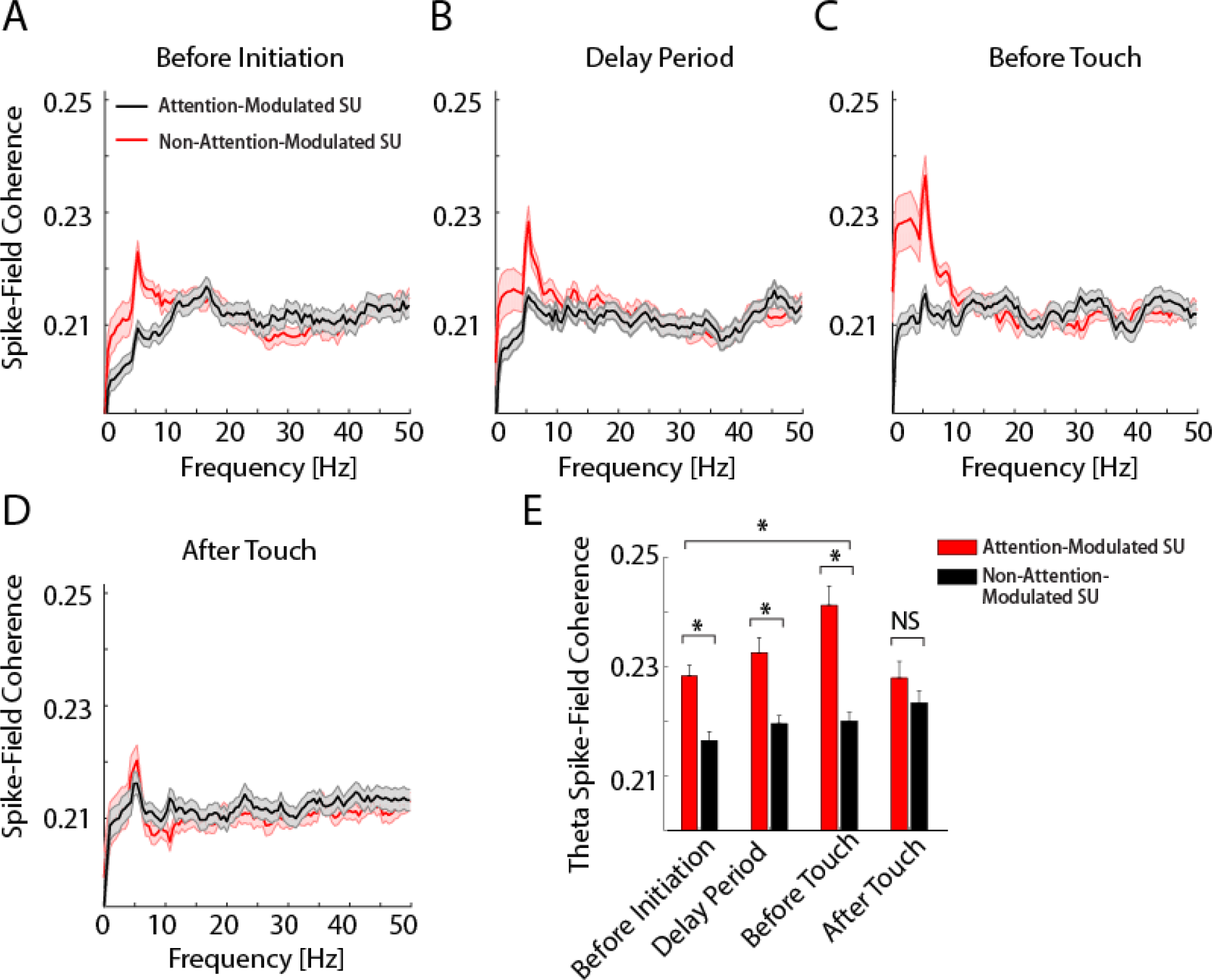
Interaction of Spiking Activity and the Local Field Potential in the Theta Band. **A-D** Spike-field coherence across sessions and animals for different task epochs for attention-modulated (red) and non-attention modulated units (black). A, before initiation; B, delay period (from initiation to stimulus onset); C, before touch (from stimulus onset to touch response); D, after touch. Shaded areas represent SEM. **E** Spike-field coherence of attention-modulated and non-attention-modulated units in theta band (4-6 Hz) for different task epochs. Theta coherence was differentially modulated by task performance. Attention-modulated units showed significantly higher coherence than non-attention-modulated units at all task epochs, except after touch. For attention-modulated units, theta coherence increased during the delay period compared to before trial initiation and reached the maximum before touch. No change in coherence occurred for non-attention-modulated units during the task. * indicates p<0.05; NS indicates p>0.05. Error bars represent SEM.

### Theta-Gamma Phase Amplitude Coupling during Attention Allocation

Since theta rhythms selectively synchronized the spiking activity of attention-modulated neurons, we next examined if the amplitude of theta rhythms was modulated in a similar way by sustained attention during the 5-CSRTT. Indeed, following trial initiation we observed an increase in theta power that remained elevated during the sustained attention period (Fig. 4A). Theta power then peaked approximately 400ms after the presentation of the visual stimulus, before falling to levels lower than before trial initiation. Theta rhythms have been observed across several cortical and subcortical brain regions, and have been implicated in the functions as diverse as working memory and spatial navigation [28]. However, these functions of theta oscillations are underpinned by one unifying mechanism: lower frequency theta oscillations temporally coordinate higher frequency gamma oscillations through phase amplitude coupling (PAC). Therefore, to test if theta oscillations in LP/Pulvinar behaved in a similar way to theta rhythms observed in other brain regions, we computed PAC between the phase of low frequency (< 10Hz) and the amplitude of high frequency LP/Pulvinar LFP signals (15-80 Hz). This analysis revealed that the amplitude of gamma oscillations (30-60 Hz) was temporally coupled to the phase of theta oscillations (4-6 Hz, Fig. 4B, PAC = 0.29 ± 0.01 SEM), confirming that LP/Pulvinar theta oscillations exhibit similar cross-frequency interaction as theta observed in other brain regions.

**Figure 4:**
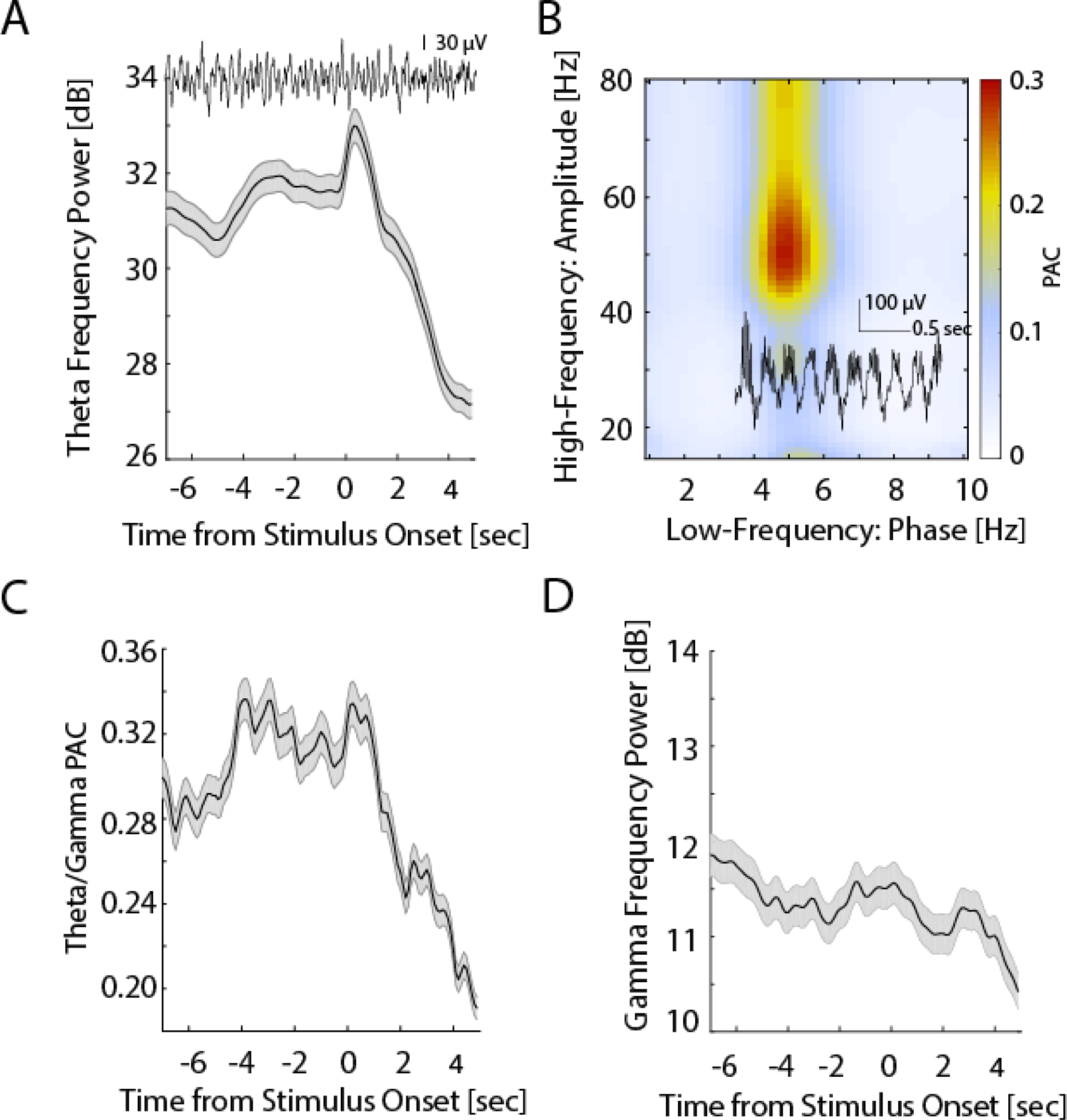
Task-Modulated Theta-Gamma Phase Amplitude Coupling (PAC) **A** Modulation of theta power during task performance. Theta power gradually increased and was maintained during the delay period and peaked between stimulus onset and screen touch. Shaded area represents SEM. **B** Heatmap of phase-amplitude coupling (PAC) during task performance shows selective coupling of theta and gamma oscillations. **C** Task-dependent modulation of time-resolved theta-gamma PAC. Theta-gamma PAC was elevated throughout the delay period and decreased after screen touch. Shaded area shows SEM. **D** Modulation of gamma power during task performance. Gamma power gradually decreased during the delay period. Shaded area represents SEM.

If theta/gamma PAC represented an underlying mechanism that LP/Pulvinar uses to selectively synchronize cortical inputs and outputs based on behavioral demands, then one would expect the strength of theta/gamma PAC to fluctuate across trials of the 5-CSRTT. To test this hypothesis, we computed theta/gamma PAC in a temporally resolved manner across the trial. We indeed found task-dependent modulation of theta/gamma PAC during performance, where PAC was significantly elevated throughout the sustained attention period, and maintained a high level until screen touch (Fig 4C, paired t-test for PAC values between delay and baseline, p < 0.001). We further examined gamma power across the trials. In contrast to the time-course of theta power and theta/gamma PAC, we found a significant reduction in gamma power during the sustained attention period (Fig 4D, paired t-test, p < 0.01). Collectively, these results indicate that even though gamma power is reduced during sustained attention, ongoing fluctuations in gamma amplitude are more tightly locked to the phase of LP/Pulvinar theta oscillations in a behaviorally dependent manner.

## Discussion

We have shown that the LP/Pulvinar complex exhibits neuronal dynamics that are modulated during the 5-CSRTT, in particular during the delay period before onset of the stimulus. We found that oscillatory activity in the theta frequency binds the neurons that were activated during the delay period. Neuronal firing of these attention-modulated neurons exhibited a ramp-like increase in their firing rate during the delay period. These results suggest that the LP/Pulvinar may play a functional role in sustained attention.

The pulvinar has only recently become a brain area of wider interest and surprisingly little is known about its function. As a major part of the visual thalamus, the pulvinar appears to play an important role in the control of visual processing and attention [29]. Two recent studies reported somewhat different findings with regards to the role of pulvinar in modulating cortical network dynamics in tasks that required selective spatial attention, which is conceptually distinct from the sustained attention investigated in our study. The work by Saalmann and colleagues [30] identified enhanced effective connectivity from pulvinar to cortical areas in the alpha frequency band for attended stimuli. In contrast, work by Zhou and colleagues [31] focused on gamma-band effective connectivity that was directed from cortex to pulvinar, however, they also found an increase in alpha effective connectivity from pulvinar to cortex. Interestingly, inactivation of the pulvinar caused not only a loss of the attentional gain in the cortical stimulus representation but an overall drop in activity below baseline for the unattended stimuli. Thus, these results support a fundamental role of the pulvinar in maintaining and perhaps guiding cortical activity in attention-demanding tasks. Together with our results, these studies suggest that while the pulvinar is involved in multiple domains of attention, the corresponding activity signatures may be distinct.

Other subunits of thalamus also appear to play a role in attention. In the context of selective attention to either visual or auditory stimuli, the medio-dorsal thalamus that forms a recurrent loop with prefrontal cortex plays a key role in amplifying PFC activity that is specific to the stimulus modality used on a given trial [32]. In addition, the nucleus reticularis thalami, which contains inhibitory neurons that provide inhibition to other thalamic areas, is recruited in the same task [33]. Although these studies have investigated other types of attention than the sustained attention probed with the 5-CSRTT they, in agreement with our results, propose a comprehensive role in allocation of processing resources.

Generators of theta oscillations have been previously described in the thalamus of animal models [34] and humans [35]. Theta rhythms in the LP/pulvinar of the cat were modulated by the state of vigilance and differed from theta oscillations recorded from hippocampus [36]. In humans, thalamic theta rhythms, in particular the anterior thalamic nucleus, have been implicated in memory formation [37]. In addition, human theta oscillations exhibit phase-amplitude coupling with higher-frequency oscillations in the range of 80 to 150 Hz [38]. Despite the differences in the frequency of the amplitude-modulated signal and the thalamic area, the parallel to our findings of theta oscillations modulating gamma oscillation is of note. In quite a different context, theta bursting in thalamus is considered to be a signature of what is called thalamocortical dysrhythmia syndrome [18]. In this model, aberrant theta oscillations (in the form of bursts) emerge due to the deinactivation of the transient, low-threshold calcium current due to deafferentation or other pathology. Together with the fact that thalamic theta oscillations are associated with decreased vigilance, the question arises why we found an increase in the theta oscillations in the delay period of the 5-CSRTT. One potential answer derives from the comparison of first-order and higher-order thalamic structures. High-order thalamic structures such as the LP/Pulvinar exhibit a substantially larger fraction of rhythmically bursting cells in the awake state [39]. Thus, rhythmic synchronized activity in higher-order thalamus could serve as a “wake-up call” to cortex due to the enhanced postsynaptic effect of such synchronized thalamic activity and thus support sustained attention

The functional characterization of thalamic networks in ferrets is in its infancy. Little is known about the connectivity of different thalamic nuclei. We recently reported neurochemical subdivision of what we referred to as the LP/Pulvinar complex [40]. By combining tracer techniques and multisite electrophysiology in the anesthetized animal we showed an agreement of structural and functional connectivity between the lateral aspect of the LP/Pulvinar complex and visual cortex. This is the thalamic location we chose for the study presented here. An additional source of confusion is the different nomenclature for seemingly similar structures across species, with different naming conventions for carnivores that include the ferret [41]. We decided to use the broader term of the LP/Pulvinar complex for extrageniculate visual thalamus to avoid dogmatic disputes of researchers across model species [42]. We argue that the structural connectivity (as previously reported by us) is the more relevant information that the specific naming scheme chosen.

To our knowledge, this is the first electrophysiological study of higher-order thalamus in sustained attention using the 5-CSRTT. However, as any scientific study, the work presented here has limitations. First, our findings on the organization of the network activity in the LP/Pulvinar complex are correlative in nature and we have not used causal circuit interrogation strategies such as optogenetics. The use of these techniques in larger-brain species such as ferrets and non-human primates is still under active development and has remained in its infancy in comparison to the investigations of the mouse brain. We argue that the synthesis of research from different model species with different levels of brain complexity substantially advances the field even if the toolsets vary between them. Nevertheless, the development of targeted causal circuit perturbations in species such as the ferret are of fundamental importance and we recently reported the first successful use of optogenetics in the awake behaving ferret [22]. Second, we have not parameterized our tasks for a more detailed dissection of the behavioral components. For example, different lengths of the delay period and presentation of distracting stimuli during the delay period are experimental manipulations that will allow the future testing of hypotheses built on the results from the study presented here. Importantly, introducing competing sensory stimuli would transform the task to one that probes selective attention [43], which differs from sustained attention. Third, and lastly, we did not investigate how the thalamic signaling modulated cortico-cortical and cortico-thalamo-cortical interactions. The goal of the current study was to delineate the role of the LP/Pulvinar complex in the ferret during sustained attention. Similar investigations with multiple electrode arrays not only in thalamus but also cortical areas that are anatomically connected will be the next step.

In conclusion, our study suggests showed that the higher-order visual thalamus is engaged during sustained attention and that oscillatory activity in the theta frequency bands organizes the neural firing of the subpopulation modulated by sustained attention. Our findings reinforce the importance of the thalamus in cognitive constructs such as attention that are often studied from a cortico-centric perspective.

## Acknowledgements

Research reported in this publication was partially supported by the National Institute of Mental Health under Award Number R01MH101547 (to F.F.). The content is solely the responsibility of the authors and does not necessarily represent the official views of the National Institutes of Health.

## Conflict of Interest

FF is founder, CSO, and majority owner of Pulvinar Neuro LLC, which markets non-invasive brain stimulation research equipment unrelated to the work presented here but has been named after the corresponding author’s favorite brain structure.

